# Dynamic Modulation of Distractor Suppression by Tonic and Trial-Level Alertness Fluctuations: A Pupillometric Study

**DOI:** 10.64898/2026.06.24.733323

**Authors:** Siyi Chen, Hermann J. Müller, Zhuanghua Shi

**Affiliations:** Ludwig-Maximilians-Universität München, 80802 München, Germany

**Keywords:** visual search, statistical learning, pupil size, reactive suppression, proactive suppression

## Abstract

Attentional control balances proactive suppression of predictable distractors with reactive suppression of unexpected ones. Yet, how internal states such as alertness shape this balance is unclear. Using pupillometry and eye tracking across two probability-cueing experiments (conducted in 2024) with varying distractor prevalence, we distinguished tonic (baseline pupil size across blocks) from trial-level pupil size fluctuations (trial-by-trial residual variability in pre-stimulus pupil size). With moderate prevalence, suppression of frequent-region distractors developed gradually, whereas high prevalence induced near-immediate suppression. Behavioral measures (e.g., reaction times) were closely linked to tonic and trial-level pupil size fluctuations. Critically, both alertness components jointly influenced control: during early learning, heightened trial-level pupil size increased distractor capture and reduced target fixations, whereas later on, suppression shifted to a proactive mode resilient to trial-level fluctuations. Under high prevalence, this shift occurred faster. Notably, higher trial-level pupil size generally accelerated first target selection. These findings show that tonic alertness and trial-level alertness fluctuations dynamically regulate reactive and proactive control during statistical learning.

**Impact Statement:** This study shows that people become better at ignoring predictable distractions over time, but that this improvement depends not only on what they have learned about the task environment, but also on their current level of alertness. By combining eye tracking and pupil measures, we found that temporary increases in alertness can sometimes help people orient more quickly to relevant information, yet during earlier stages of learning they can also make attention more vulnerable to distracting events. These findings suggest that successful focus in complex environments depends on a dynamic interplay between learned expectations and moment-to-moment fluctuations in mental state, with implications for understanding sustained attention in settings such as monitoring, driving, and other tasks that require people to stay engaged while resisting distraction.

## Introduction

Visual attention operates through a delicate balance between enhancing goal-relevant ‘target’ information and suppressing potentially irrelevant ‘distractor’ stimuli. Crucially, distractor suppression operates in two complementary modes: proactive and reactive. Proactive suppression is anticipatory, attempting to prevent attentional capture before it occurs; reactive suppression reallocates attention to relevant targets following capture by a distractor (Liesefeld et al., 2024; Sauter et al., 2021; Geng, 2014). Proactive strategies are efficient but depend on predictability; reactive strategies are resource-demanding, involving a chain of recovery processes.

A growing body of work shows that proactive distractor suppression is shaped by statistical learning. Regularities such as distractor prevalence, location, or feature predictability allow proactive control to minimize capture, while unpredictable distractor events force greater reliance on reactive suppression (Chen et al., 2024; Chen et al., 2025; Failing & Theeuwes, 2018; Gaspar & McDonald, 2014; Gaspelin & Luck, 2018; Geng & Behrmann, 2005; Sauter et al., 2019; Zhang et al., 2019, 2022). For instance, high distractor prevalence promotes proactive suppression, whereas low prevalence invokes trial-by-trial reactive control (Müller et al., 2009; Vatterott & Vecera, 2012; Won et al., 2020). When distractors repeatedly occur in a specific region, statistical learning enables observers to anticipate their location, making them less likely to capture eye movements and reducing interference with performance (Allenmark et al., 2024; Sauter et al., 2021).

Dealing with distraction consumes cognitive resources, and the availability of these can vary across time and individuals, so that distractors sometimes capture attention even when capture could, in principle, be avoided. A central source of variability in attention is vigilance: the ability to sustain focus and readiness over prolonged, often monotonous periods of task performance (Oken et al., 2006). Conceptually, ‘vigilance’ overlaps with intrinsic or tonic alertness, a self-generated baseline state independent of external cues. Trial-level alertness fluctuations, by contrast, refer to moment-to-moment changes in readiness that arise from internal state variability and recent task events (Langner & Eickhoff, 2013; Posner, 2008; Sturm & Willmes, 2001). In the present study, we use ‘vigilance’ as an umbrella term encompassing both tonic alertness—slow, sustained fluctuations in baseline arousal—and trial-level alertness fluctuations that capture rapid, trial-by-trial variability. Classic vigilance research established that sustained attention degrades over time—the vigilance decrement—and that arousal level critically modulates perceptual sensitivity and response readiness (Broadbent, 2013; Posner, 1978). Yet, despite these well-documented links between arousal and performance, contemporary accounts of statistical distractor learning have not examined how tonic and trial-level alertness states shape the balance between reactive and proactive suppression as it unfolds during learning. This gap is consequential, because alertness fluctuations generate specific predictions: heightened arousal should benefit reactive suppression by increasing the resources available for recovery from capture, but may simultaneously disrupt the fine-tuned proactive control that depends on stable, automatized task sets; moreover, the transition from reactive to proactive suppression should depend on both learning opportunity (i.e., distractor prevalence) and the observer’s prevailing arousal state. Indeed, proactive and reactive suppression may differ in their reliance on vigilance. Early on during learning, before distractor regularities are consolidated, suppression likely requires active control rather than operating automatically (Shiffrin & Schneider, 1977)—leaving a window for distraction. If a distractor happened to capture attention, reactively suppressing it is inherently resource-demanding: the distractor must be identified as a non-target item and attention disengaged from it to reimpose goal-directed attentional orienting (Sauter et al., 2021). These processing costs may be minimized later on, once suppression has become largely automatized, operating proactively (Sauter et al., 2021; Zhang et al., 2022). But even the latter mode requires the maintenance of task-appropriate control settings—that is, an active task set—within and across trials (Allenmark et al., 2024; Qiu et al., 2024; Zhang et al., 2024).

Thus, despite recent advances in understanding the mechanisms that underlie statistical distractor learning, a critical gap remains: how do trial-level alertness fluctuations, as well as more tonic, slower-changing states of vigilance, influence the acquisition and the use of learnt attentional priorities? This question is particularly important as statistical learning unfolds over time, requiring sustained engagement to extract environmental regularities, consolidate them into automatized routines, and call upon these as demanded by the task. Addressing this gap requires a framework that connects vigilance with the underlying neurocognitive mechanisms. The adaptive-gain theory (Aston-Jones & Cohen, 2005) provides a useful conceptual backdrop by linking locus coeruleus–norepinephrine (LC–NE) dynamics to sustained and transient alertness, distinguishing between two modes of activity: a phasic mode, marked by burst-like responses that facilitate exploitation of task-relevant information, and a tonic mode, characterized by sustained activity that promotes disengagement and exploration when task utility declines. Importantly, tonic LC activity exhibits an inverted-U relationship with attentional performance, mirroring the Yerkes–Dodson law (Yerkes & Dodson, 1908; Hockey, 1997): very low activity is linked to drowsiness, very high activity to distractibility, while moderate activity—when coupled with strong phasic responses to goal-relevant stimuli—supports optimal performance (see also Gilzenrat et al., 2010; Minzenberg et al., 2008). While pupil dynamics have been linked to LC-NE function (Beatty, 1982; Joshi & Gold, 2020; Mathôt, 2018), it is important to note that pupil size reflects the combined influence of multiple neuromodulatory and autonomic systems, and the mapping between pupil measures and specific LC firing modes is indirect. Specifically, baseline pupil size has been associated with tonic LC activity, where intermediate levels are associated with optimal task engagement (Gilzenrat et al., 2010; Jepma & Nieuwenhuis, 2011; van den Brink et al., 2016). In contrast, task-evoked pupil dilations have been linked to phasic LC bursts that support correct detections and adaptive decision-making (de Gee et al., 2021; Murphy et al., 2011; Reimer et al., 2014; van den Brink et al., 2016). While prior work has often examined baseline pupil size or trial-related responses separately, few studies have disentangled slow, block-wise trends from rapid fluctuations in alertness. Here, we model baseline pupil size across blocks to index tonic alertness and use residual variability to capture trial-level alertness fluctuations, providing an empirical account of how alertness dynamics relate to the calibration of attentional control.

Thus, in the present study, we investigated whether tonic and trial-level alertness fluctuations shape distractor-handling mechanisms—specifically, reliance on proactive versus reactive control. Drawing on the open dataset of Allenmark et al. (2024), which, within and across two experiments, varied distractor region and distractor prevalence, we examined how statistical learning and vigilance interacted over the course of prolonged experimental sessions. Focusing on first-fixation behavior as an early measure of attentional capture, we hypothesized that in Experiment 1 (50% distractor prevalence), distractor costs would be largest during early learning when tonic alertness is high and trial-by-trial alertness fluctuations strongly influence performance, requiring reactive control, which could be indicated by a high rate of distractor-driven oculomotor capture. As learning progresses and suppression becomes proactive, evidenced by a declining capture rate, trial-level alertness effects should diminish along with the decline in tonic alertness. In Experiment 2 (with distractor prevalence increased to 70%), we expected proactive suppression to develop faster, reducing trial-level alertness effects on distractor capture.

## Method

### Participants

The present analyses are based on the existing dataset of Allenmark et al. (2024), with sample sizes of N = 15 (7 females; Mean age = 28.3, SD = 3.2 years; Experiment 1) and N = 16 (9 females; Mean age = 26.6, SD = 4.2; Experiment 2). These samples were determined by the original study design, which was powered for the primary behavioral probability-cueing effects—confirmed here with >99% power. For the within-experiment analyses reported here, post-hoc power calculations for the significant effects confirmed that all exceeded 85% power. Specifically, the large number of trials contributed by each participant (∼1008 observations), allowing reliable estimation of within-participant trial-level relationships between pupil-linked state and gaze behavior.

### Experiment Design and Procedure

In the study of Allenmark et al. (2024) participants searched for two tilted target bars (“i”-type; ±12°) among vertical non-target bars (“i”-type). On distractor-present trials, one vertical bar was replaced by a salient horizontal bar (**Figure 1**), which appeared with high probability (90%) in one half of the display (top or bottom, counterbalanced across participants) and with low probability (10%) in the opposite half, thereby creating a probability-cueing manipulation. This manipulation defined the factor Distractor Region, which comprised three levels: Frequent-Region Distractor (distractor bar at high-probability location), Rare-Region Distractor (distractor at low-probability location), and Distractor Absent. The two targets had an equal chance of appearing at any of the 10 (used) locations on the intermediate ring, but never at directly adjacent locations and (on distractor-present trials) never at the distractor location.

**Figure 1.**
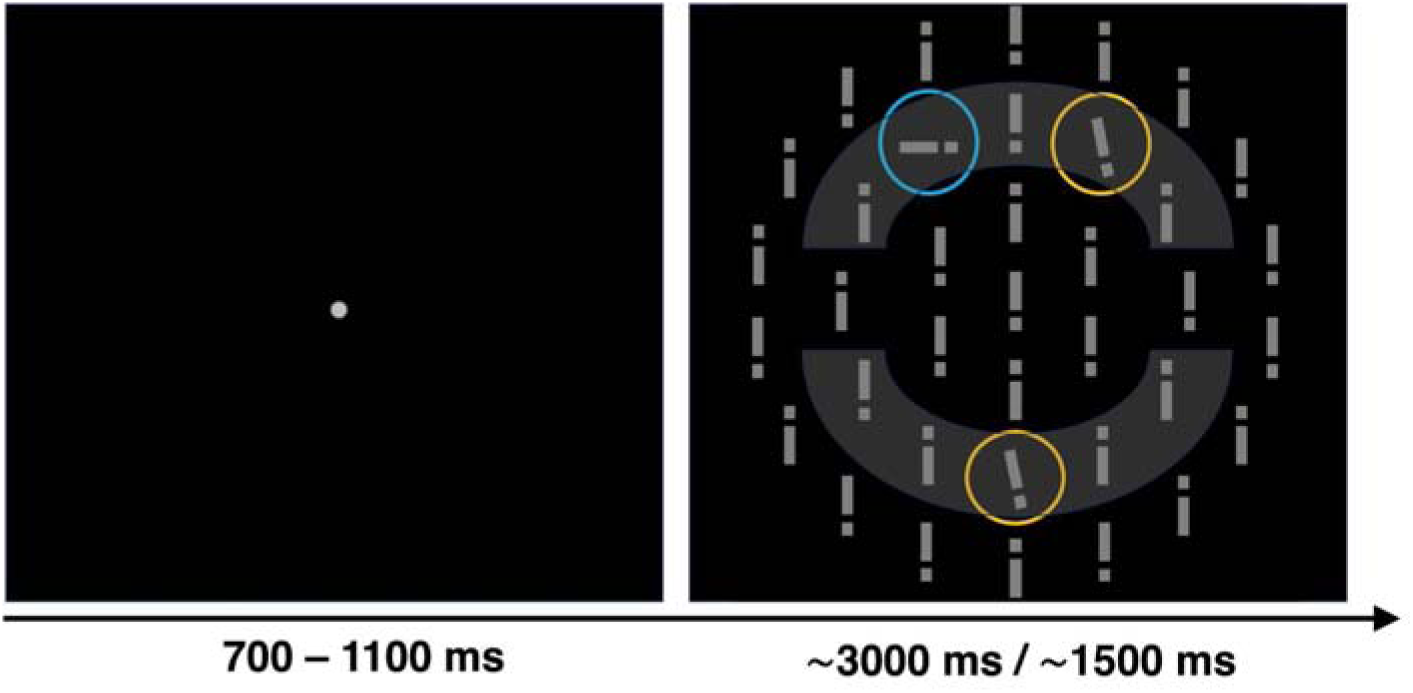
Trial sequence and search display (adapted from Allenmark et al., 2024). Each trial began with a central fixation dot (700–1100 ms), followed by a search display. Participants had to fixate the central dot and then, upon exposure to the search display, move their eyes towards the two slightly tilted target bars (±12°) in any order. Their task was to identify the positions of the small “i” dots on both targets and make a speeded 2AFC-Go response versus a No-Go response: if both dots were at the top or the bottom, they had to make a “top” vs. “bottom” decision by pressing either the “J” or the “F” key on the keyboard (“Go” trial; right panel); if the positions of the dots differed, they withheld their response (“No-Go” trial). The search display remained on until response (max. 3000 ms, Experiment 1) or, respectively, for 1500 ms followed by a mask (1500 ms) replacing the ‘i’s by ‘I’s (Experiment 2). The gray ellipses mark the frequent and rare distractor regions; yellow dotted circles indicate target bars, and the blue circle indicates the salient distractor (horizontal bar). Color overlays are for illustration only.

Participants were instructed to fixate the targets and, on Go trials (when i-dots of both targets were in the same position, top or bottom), report the dot location. In contrast, on No-Go trials (when the targets’ i-dots were in different positions), they withheld their response. In Experiment 1, distractors appeared on 50% of trials, the search display remained visible until response (max. 3000 ms), and 20% of trials were No-Go trials. In Experiment 2, distractor prevalence increased to 70%, the display was visible for 1500 ms before a 1500 ms mask removed the response-critical dot-position information, and 40% of trials were No-Go trials.

In this task, errors could occur in various ways. First, participants could misjudge the trial type: if the two i-dot positions were different, the correct response would have been No-Go, so making a Go response on a No-Go trial counted as an error; and vice versa for Go trials. Second, even on Go trials, in which the two i-dot positions were the same (and an overt response was in principle correct), participants could still make an error by misclassifying the i-dot position, for example, responding “top” when the correct answer was “bottom”. Note that participants were not given error feedback upon their response decisions at the end of a trial.

Each experiment consisted of 1008 trials, arranged in 24 blocks, which took approximately 90 minutes to complete. Participants were free to take breaks of a self-determined length between consecutive blocks.

### Recording, Pupil Preprocessing, and Trial Exclusion

Participants viewed the stimuli presented on a 21-inch SONY GDM-F500R CRT monitor (120 Hz, 1024 × 768), with their heads supported by a chin rest and forehead support. Eye movements and pupil diameter were recorded throughout the task with an SR Research EyeLink 1000 system (1-kHz sampling; default saccade criteria: 35°/s velocity, 9500°/s² acceleration). Raw pupil traces were corrected for blink-related artifacts by linearly interpolating samples within ±150 ms of detected blinks, low-pass filtered with a third-order Butterworth filter at 10 Hz, and cleaned of residual blink- and saccade-related transients with pupil impulse response deconvolution (Hoeks & Levelt, 1993). Trials with response errors, excessive blinks (>20%), missing data, fixation durations below 50 ms, or saccade latencies below 80 ms were excluded from eye-movement analyses, removing 6% of trials. For more detailed information about the original task and recording setup, see Allenmark et al. (2024).

### Statistical Analysis

#### Bayesian Learning Models for Initial Fixations

After preprocessing the eye and pupil data, we first tracked statistical learning with Bayesian mixed-effects logistic-regression models (brms; Bürkner, 2017) predicting the probability of first fixation from time-on-task (log-transformed block number), distractor region (frequent, rare, distractor-absent), and their interaction, including per-participant random intercepts and slopes:

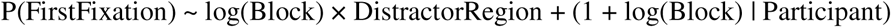

In accordance with prior research demonstrating robust probability-cueing effects regardless of response type (Allenmark et al., 2024), Go and No-Go trials were collapsed for the statistical learning analyses. Initial fixations were coded as two binary outcomes: first fixation on a target (1 vs. 0) and first fixation on the distractor (1 vs. 0). Target-fixation models included frequent-region distractor, rare-region distractor, and distractor-absent trials. Distractor-fixation models were restricted to distractor-present trials because a distractor fixation is structurally impossible when no distractor appeared. These outcomes served as indices of early attentional selection and distractor capture.

#### Model Reporting and Diagnostics

All primary mixed-effects analyses were fitted in brms as Bayesian models. Binary fixation and error outcomes used Bernoulli-logit models, whereas continuous RT and latency outcomes used Gaussian models. Models used weakly informative priors for fixed effects (Normal [0,1]) and group-level standard deviations (Student-t [3, 0, 2.5]), with four MCMC chains per model (4000 iterations, 2000 warmup). Convergence was assessed using R^ (< 1.01), effective sample sizes, divergent transitions, and maximum-treedepth warnings. Bayesian effects are summarized in terms of posterior means, 95% credible intervals, and posterior probabilities.

#### Baseline-Pupil Decomposition

Baseline pupil size was computed as the mean pupil diameter during the 500 ms preceding stimulus onset for each trial, excluding missing samples and z-scoring values within participants. This pre-stimulus window reduced contamination from current-display luminance and stimulus-evoked pupil responses. To separate slow block-wise drift from trial-specific variation, we fitted a participant-level regression of baseline pupil size on log(Block). The fitted values captured the slow tonic-pupil trend and were used to define pupil-linked task phases. By construction, this measure is coupled with time-on-task and learning. The residuals captured trial-level deviations around the fitted trend and substantially reduced smooth block dependence, providing the primary trial-level pupil measure for testing state-dependent selection effects. Analyses were restricted to pre-stimulus fixation periods, values outside ±3 SD were excluded, and task-evoked pupil responses were analyzed separately as a check (see supplementary “Stimulus-Triggered Pupil Response Analysis”). The baseline-trend regression, phase-validation ANOVA, and recent-trial ANOVAs were treated as descriptive checks supporting the pupil decomposition; the focal inferential tests used the Bayesian mixed-effects models described below.

#### Reaction-Time Models

We tested whether pupil-linked measures predicted behavioral performance by fitting Bayesian Gaussian mixed-effects models to RTs on correct Go trials, excluding RTs below 200 ms and RTs more than 3 SD from each participant’s mean within each distractor-region condition. The corresponding model for error responses is reported in the Supplemental Materials (see supplementary “Tonic and Phasic Pupil Dynamics in Error Responses”).

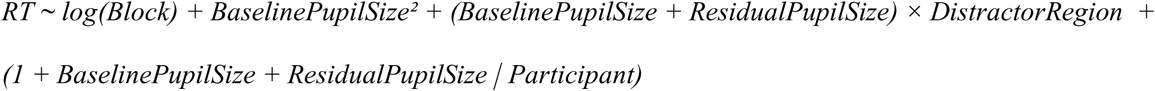

These fixed effects separated slow tonic-pupil variation from trial-level residual pupil fluctuations. Baseline pupil size and its quadratic term captured potentially non-linear associations between tonic pre-stimulus pupil level and RT (Yerkes & Dodson, 1908; Aston-Jones & Cohen, 2005), whereas log(Block) adjusted for practice, fatigue, and learning-related changes across the task. Interactions with Distractor Region tested whether pupil-linked state related differently to performance in frequent, rare, and absent distractor-region trials. Residual pupil size indexed momentary trial-to-trial deviations from the slow pupil trend. Participant-level random intercepts and random slopes for baseline pupil size and residual pupil size allowed observers to differ in overall RT and in the strength of pupil–RT coupling, with estimates partially pooled across participants.

#### Learning-Phase-Dependent Residual-Pupil Effects

To examine whether trial-level pupil effects changed across the slow pupil-defined phases of the task, we fitted block-adjusted Bayesian mixed-effects models for first-fixation probability and first-saccade latency. The fixation model used a Bernoulli-logit likelihood, whereas the latency model used a Gaussian likelihood; both included residual pupil size, tonic-pupil phase, distractor region, and their interaction, together with continuous block-related learning terms:

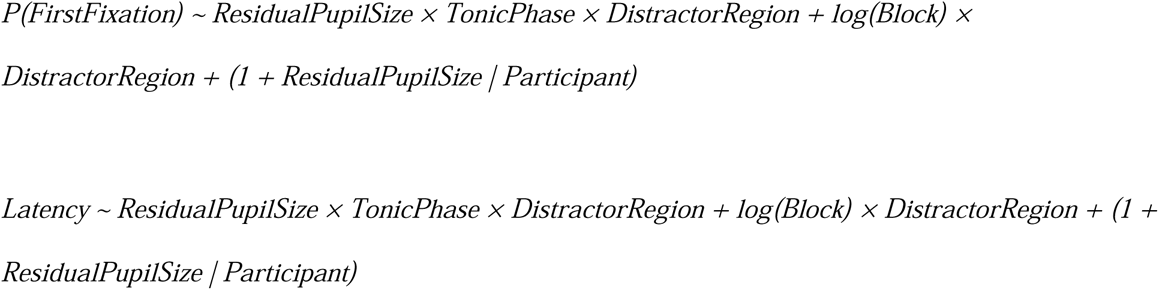

Because residual pupil size was the focal trial-level predictor, participant-level random slopes were included for residual pupil size to allow pupil–behavior coupling to vary across observers. We used this targeted random-effects structure rather than a fully maximal structure to keep the model estimable given the sample size and the number of binary fixation events in some cells.

We derived tonic-pupil phases (high, intermediate, low) from fitted baseline pupil trends (BaselinePupilSize(z) ∼ log(Block)). Trials with fitted values ≥ 0.1 were classified as high, ≤ –0.3 as low, and the remaining trials as intermediate. These thresholds were chosen to create descriptively separated pupil-linked phases while retaining a sufficient and roughly comparable number of trials in each phase for mixed-effects modeling (trial counts in Exp.1: M(high) = 333 [305, 361]; M(intermediate) = 338 [248, 429]; M(low) = 387 [342, 431]; Exp.2: M(high) = 257 [204, 310]; M(intermediate) = 499 [360, 638]; M(low) = 334 [287, 381]). The derived tonic-pupil phases differed strongly in raw baseline pupil size (Experiment 1: *F*(1.53, 18.39) = 36.44, *p* < .001, η_p_^2^ = .75; Experiment 2: *F*(1.44, 14.39) = 37.17, *p* < .001, η_p_^2^ = .79): the baseline pupil size was larger at high tonic level (Exp.1: M = 0.68; Exp.2: M = 0.65) compared to both intermediate (Exp.1: M = -0.14, p = .0004; Exp.2: M = -0.16, p < .001) and low levels (Exp.1: M = -0.5, p < .001; Exp.2: M = -0.38, p < .001). Because these phases are derived from the block-wise trend, they are not intended to separate tonic pupil level from time-on-task. The log(Block) × DistractorRegion term accounts for condition-specific learning trajectories, allowing the model to ask whether residual trial-level pupil fluctuations have different behavioral consequences across tonic-pupil phases beyond block-related changes in distractor handling. This model tests whether the trial-level residual-pupil slope differs across pupil-linked task phases after adjusting for region-specific learning.

For interpretation, we defined the evidence hierarchy in advance of the Results: first fixations on the distractor were the central test of oculomotor capture; first fixations on the target were treated as complementary support; and saccade-latency effects were interpreted as readiness or response-vigor effects.

#### Complementary Fixation-Stage Analysis

As a complementary temporal-localization analysis, we used the two-target structure of the task to classify distractor fixations by when they occurred in the eye-movement sequence. Following the logic of Allenmark et al. (2024), distractor fixations were coded as occurring before the first target was fixated, between the first and second target fixations, or after both targets had been inspected. These fixation-stage Bayesian logistic models used the same residual-pupil and block-adjustment structure as the first-fixation models. Because post-target fixations are response-dependent in this paradigm, the post-target model additionally included Response × DistractorRegion.

## Results

### Learning Profiles Differed Across Experiments

To examine how participants learned to suppress distractors during early attentional processing, we analyzed first-fixation behavior across blocks using a Bayesian mixed-effects logistic model (P(FirstFixation) ∼ log(Block) × DistractorRegion + (1 + log(Block) | Participant)).

In Experiment 1 (50% distractor prevalence, **Figure 2a**), distractor capture in the frequent region decreased with increasing log(Block) (β = -7%, 95% CrI [-10%, -4%], Pr(β < 0) > .999), and the log(Block) × Distractor Region interaction showed that the rare--frequent difference increased across blocks (β = 9%, 95% CrI [4%, 13%], Pr(β > 0) > .999). Target fixations showed the complementary pattern, increasing with log(Block) in the frequent condition (β = 7%, 95% CrI [4%, 10%], Pr(β > 0) > .999), while remaining stable in the rare and distractor-absent conditions (rare-region distractor: β = -1%, 95% CrI [-4%, 2%], Pr(β < 0) = .74; distractor-absent: β = 1%, 95% CrI [-2%, 4%], Pr(β > 0) = .74). These results, in particular the interaction effects, provide direct evidence that distractor regularities were gradually learned.

**Figure 2.**
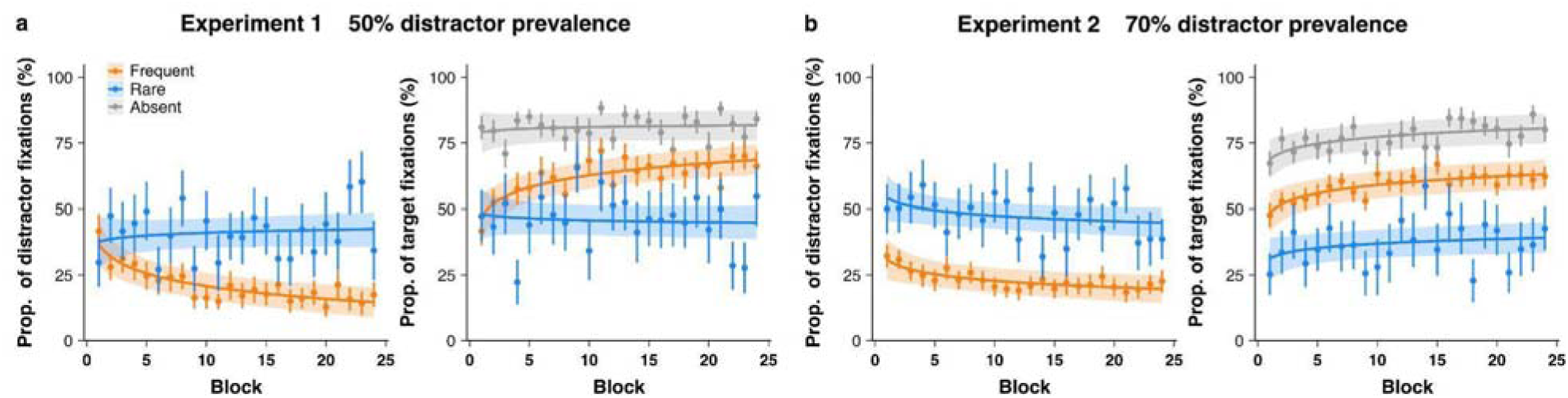
Predicted and observed proportion of distractor and target fixations across blocks (42 trials per block) as a function of distractor location (frequent region = orange, rare region = blue), presented separately for Experiment 1 (a) and Experiment 2 (b). Curves show the Bayesian model predictions (with 95% credible intervals shaded) from a model (P(FirstFixation) ∼ log(Block) × DistractorRegion + (1 + log(Block) | Participant)). Points show empirical mean fixation proportions per block, with error bars representing ±1 standard error (SE).

In Experiment 2 (70% distractor prevalence, **Figure 2b**), learning effects emerged earlier. Distractor capture in the frequent region decreased modestly with increasing log(Block) (β = -4%, 95% CrI [-6%, -1%], Pr(β < 0) = .999), but the log(Block) × Distractor Region interaction was absent (β = 1%, 95% CrI [-3%, 5%], Pr(β > 0) = .69); participants already differentiated distractor regions, showing 23% less capture by frequent- than rare-region distractors from the outset (95% CrI [13%, 33%], Pr(β) > .999). Target fixations similarly increased with log(Block) (β = 5%, 95% CrI [2%, 7%], Pr(β > 0) > .999) and were less likely for rare- than frequent-region distractors (β = -17%, 95% CrI [-27%, -8%], Pr(β < 0) > .999). These results indicate that in the Experiment 2 task context, efficient suppression for frequent-region distractors was present early, with participants showing reduced capture by frequent-region distractors even from the first block.

To formally test whether learning dynamics differed between experiments, we combined both datasets and fitted a trial-level Bayesian logistic mixed model for distractor-present trials. The model predicted first fixation on the distractor from Experiment, Distractor Region, log(Block), and their interactions, with participant-level random intercepts and log(Block) slopes. The focal Experiment × Distractor Region × log(Block) interaction was credibly negative (β = -0.366 log-odds, 95% CrI [-0.614, -0.124], Pr(β < 0) > .999), showing that the rare-minus-frequent learning-slope contrast was smaller in Experiment 2 than in Experiment 1. We therefore used the cross-experiment model to characterize the datasets as differing in learning trajectory, providing the learning-phase context for the pupillometric analyses.

### Baseline Pupil Dynamics and Recent Trial Events

Having established statistical learning, we next decomposed baseline pupil size into a slow block-wise trend and trial-by-trial residual fluctuations. Baseline pupil size declined reliably with log(Block), consistent with a gradual reduction in tonic-pupil level over task progression. This decline was significant in both experiments, though steeper in Experiment 1, β = -0.636, SE = 0.009, 95% CI [-0.653, -0.619], *p* < .001, than in Experiment 2, β = -0.458, SE = 0.009, 95% CI [-0.476, -0.441], *p* < .001; the Experiment 2 × log(Block) interaction confirmed the shallower decline in Experiment 2, β = 0.177, SE = 0.012, *p* < .001. We interpret this fitted trend as a tonic-pupil phase marker that tracks the slow task progression. The standardized residuals captured trial-level deviations around this trend and varied considerably within blocks. **Figure 3** shows block-wise residual means for visualization in panels **a** and **b** and representative trial-level residual sequences in panels **c** and **d**; the statistical analyses use each participant’s trial-level residual values (see **Methods**).

**Figure 3.**
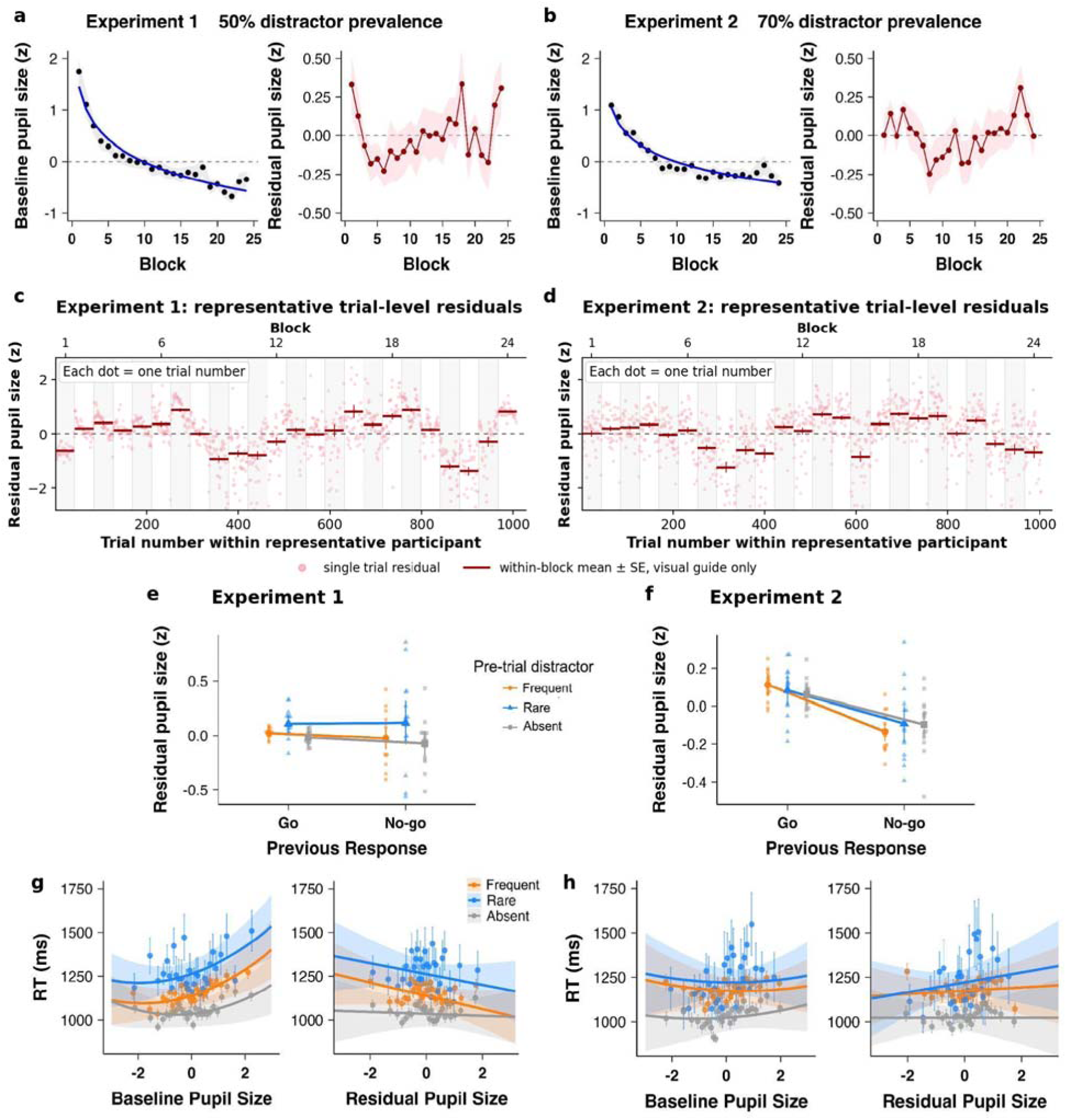
Baseline pupil dynamics and trial-level residual pupil fluctuations. Panels **a** and **b** show, for Experiments 1 and 2 respectively, the block-wise group trajectory of baseline pupil size and the residuals after removing the log-block trend. All pupil measures were z-scored within participants. Black points show observed block means across participants, the blue curve shows the fitted trend from the model BaselinePupilSize(z) ∼ log(Block), and red points show block-wise means of standardized residuals. Shaded ribbons indicate ±1 SEM. Panels **c** and **d** show trial-level residual pupil values from one representative participant in each experiment. Each pink point is a single trial plotted at its recorded trial number; block labels and vertical separators are shown on the nominal 42-trial block structure and are provided only to orient the sequence. Red horizontal segments show within-block mean ± SE as a visual guide and were not the model input. Panels **e** and **f** summarize pre-stimulus residual pupil size on the current trial as a function of the previous trial’s response type (Go, No-Go) and distractor region (Frequent, Rare, Absent). Panels **g** and **h** provide exploratory descriptive visualizations of Go-trial RTs as a function of baseline pupil size and trial-level residual pupil size, separately by distractor condition. Points in panels **g** and **h** are binned averages (30 bins per condition), lines show descriptive fitted trends, and error bars indicate ±1 SE.

To investigate whether residual pre-stimulus pupil size was related to recent task events, we conducted a within-subject ANOVA with previous-trial distractor region and previous-trial response type. In *Experiment 1* (50% distractor prevalence; **Figure 3e**), residual pupil size was higher following rare-region distractors than following frequent-region or distractor-absent trials (*F*(1.15, 16.14) = 4.93, *p* = .037, *η_p_^2^* = .26). This effect did not depend on whether the rare-region distractor captured the eye (*p* = .58, *η_p_^2^* = .02). In *Experiment 2* (70% prevalence; **Figure 3f**), residual pupil size was instead higher following Go than No-Go trials (*F*(1, 15) = 18.57, *p* < .001, *η_p_^2^* = .55). Thus, residual pupil size was not random noise around the slow baseline trend; it also reflected recent task events.

A Bayesian mixed-effects model was used to predict reaction times (RTs) on Go trials from baseline pupil size, baseline pupil size squared, residual pupil size, distractor region (reference = frequent-region distractors), their interactions, and log(Block). **Figure 3g** and **Figure 3h** depict these effects for Experiments 1 and 2, respectively.

In *Experiment 1*, RTs decreased over blocks (β = -61.03 ms, 95% CrI [-76.29, -45.54], Pr(β < 0) > .999). Larger baseline pupil size predicted slower RTs (β = 55.26 ms, 95% CrI [31.47, 78.92], Pr(β > 0) > .999), with an additional positive quadratic effect (β = 8.20 ms, 95% CrI [4.43, 12.00], Pr(β > 0) > .999). Distractor-region effects were also robust: RTs were slower for rare- than frequent-region distractor trials (β = 119.72 ms, 95% CrI [93.02, 147.14], Pr(β > 0) > .999) and faster for distractor-absent than frequent-region distractor trials (β = -105.62 ms, 95% CrI [-117.82, -93.28], Pr(β < 0) > .999). Critically, larger residual pre-stimulus pupil size predicted faster RTs in frequent-region distractor trials (β = -45.98 ms, 95% CrI [-72.05, -19.93], Pr(β < 0) > .999). Simple slopes showed the same negative direction for rare-region trials (β = -34.58 ms, 95% CrI [-86.91, 19.19], Pr(β < 0) = .893) and distractor-absent trials (β = -13.19 ms, 95% CrI [-39.00, 13.31], Pr(β < 0) = .842), but these latter intervals included zero.

In *Experiment 2*, RTs again decreased over blocks (β = -76.65 ms, 95% CrI [-91.93, -61.70], Pr(β < 0) > .999) and showed a reliable positive quadratic baseline-pupil effect (β = 7.35 ms, 95% CrI [3.72, 10.90], Pr(β > 0) > .999), whereas the linear baseline-pupil effect was uncertain (β = -11.34 ms, 95% CrI [-42.78, 20.31], Pr(β < 0) = .760). Distractor-region effects were again evident: RTs were slower for rare- than frequent-region distractor trials (β = 44.59 ms, 95% CrI [12.40, 76.99], Pr(β > 0) = .996) and faster for distractor-absent than frequent-region distractor trials (β = -150.24 ms, 95% CrI [-164.12, -136.16], Pr(β < 0) > .999). In contrast to Experiment 1, residual pupil size did not show a robust RT effect in frequent-region trials (β = 14.94 ms, 95% CrI [-18.57, 48.84], Pr(β > 0) = .811), nor did its interactions with rare-region (β = 31.16 ms, 95% CrI [-44.87, 108.47], Pr(β > 0) = .792) or distractor-absent trials (β = -7.57 ms, 95% CrI [-42.95, 27.96], Pr(β < 0) = .663). Thus, Experiment 1 showed a clear residual-pupil/faster-RT relation, while Experiment 2 did not show a robust residual-pupil RT effect, and both experiments showed reliable block, quadratic baseline-pupil, and distractor-region effects.

### Residual Pupil Effects on Oculomotor Capture

We next examined whether residual trial-level pupil fluctuations related differently to attentional selection across tonic-pupil phases (high, intermediate, low). These phases track the slow fitted baseline-pupil trend and therefore also index task progression (see Method). The inferential models included log(Block) × DistractorRegion to adjust for condition-specific learning trajectories: P(FirstFixation) ∼ ResidualPupilSize × TonicPhase × DistractorRegion + log(Block) × DistractorRegion + (1 + ResidualPupilSize | Participant). The critical test was the residual-pupil slope for first fixations on the distractor, because this is the most direct fixation outcome for distractor capture. First fixations on the target and saccade latencies with the same model structure were used to evaluate whether the same residual-pupil fluctuations showed converging effects on selection and readiness.

#### Experiment 1: First distractor fixation

The key distractor-capture result was the residual-pupil slope for first fixations on the distractor in frequent-region distractor trials during the high tonic-pupil phase (**Figure 4a**). In this condition, higher residual pupil size predicted a greater probability of first fixating the distractor, β = 0.175, 95% CrI [0.045, 0.310], Pr(β > 0) = .996. This effect was not evident in the intermediate phase, β = 0.043, 95% CrI [-0.098, 0.185], Pr(β > 0) = .723, or low phase, β = -0.019, 95% CrI [-0.153, 0.113], Pr(β > 0) = .391. Compared with the high tonic-pupil phase, the residual-pupil slope for frequent-region distractor fixations was credibly weaker in the low phase, β = −0.194 [−0.352, −0.036], and directionally weaker in the intermediate phase, β = −0.132 [−0.300, 0.035]. The intermediate and low phases did not differ clearly, β = 0.062 [−0.108, 0.231]. The overall distractor fixation probabilities were lower in both intermediate and low phases than in the high phase (intermediate: −5.5% [−9.0%, −2.4%], β = −0.357 [−0.537, −0.171]; low: −4.3% [−8.0%, −1.0%], β = −0.272 [−0.475, −0.064]), with no credible difference between intermediate and low phases. Thus, even after accounting for condition-specific learning over blocks, residual-pupil fluctuations were associated with increased distractor capture specifically during the high tonic-pupil phase in Experiment 1. No comparable residual-pupil effects were observed for rare-region distractor trials.

**Figure 4.**
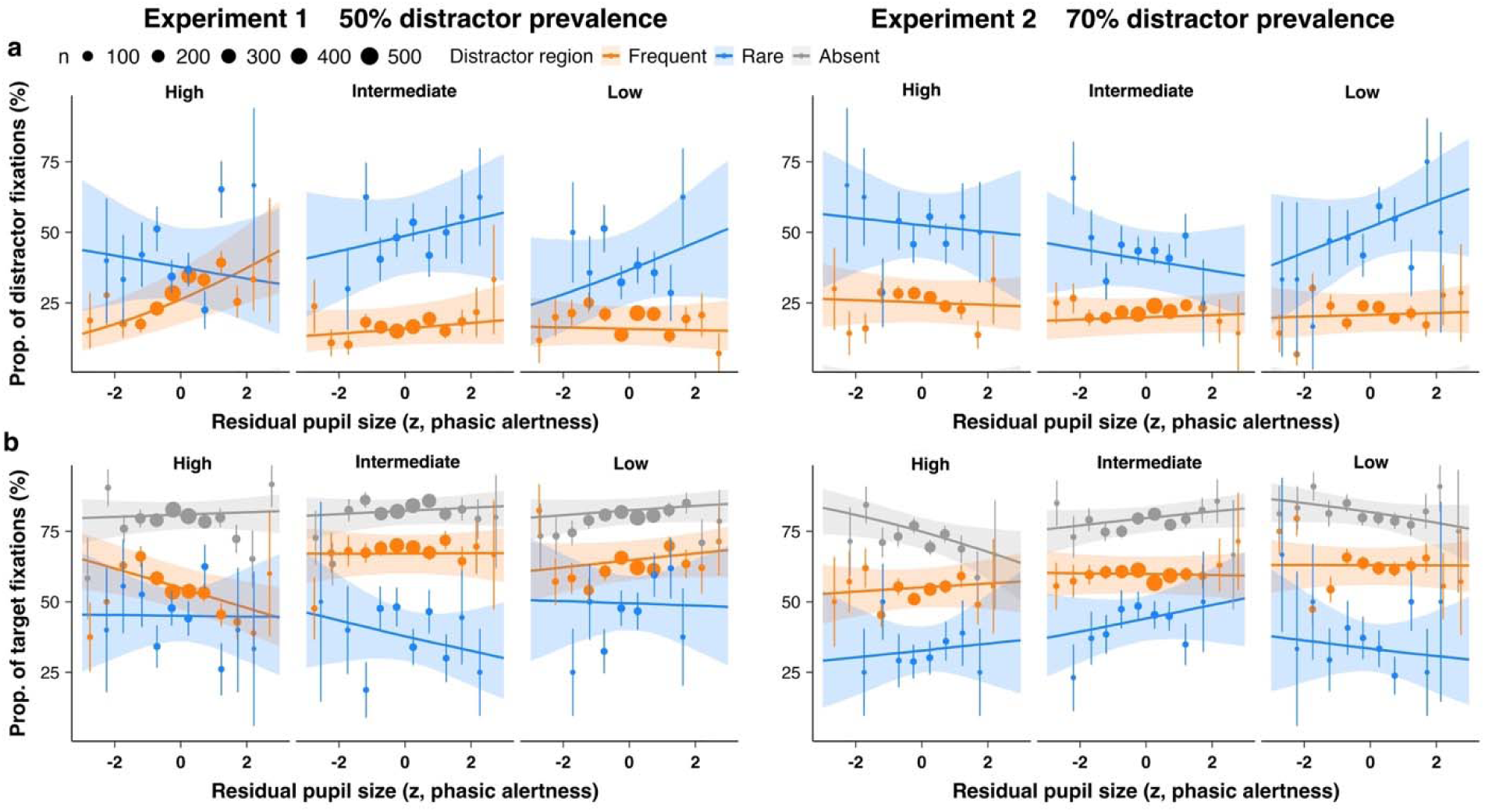
Model-predicted first-fixation probabilities from the Bayesian block-adjusted phase models. Panel (**a**) shows distractor-directed first fixations, and Panel (**b**) shows target-directed first fixations, as a function of residual pre-stimulus pupil size, tonic-pupil phase, distractor region, and experiment. Solid lines depict model-predicted trends; shaded areas depict 95% credible intervals, and points represent binned raw data with the size indicating the trial number in each condition. Error bars indicate ±1 SE.

#### Experiment 1: First target fixation

A complementary target-fixation analysis showed an opposite pattern in the same frequent-region/high-phase condition (**Figure 4b**). Higher residual pupil size predicted a lower probability of first fixating the target, β = −0.092, 95% CrI [−0.190, 0.005], Pr(β < 0) = .968. This pattern was not evident in the intermediate phase, β = 0.003, 95% CrI [−0.097, 0.102], Pr(β < 0) = .473, or low phase, β = 0.064, 95% CrI [−0.023, 0.150], Pr(β < 0) = .072. Post-hoc slope contrasts indicated that the target-fixation slope was credibly less negative in the low than high phase, β = 0.156, 95% CrI [0.030, 0.284], whereas the intermediate–high contrast was directional but uncertain, β = 0.095, 95% CrI [−0.039, 0.231]. No comparable residual-pupil effects were observed for rare-region distractor and distractor-absent trials, where all credible intervals included zero. Thus, the fixation results indicate that residual pupil fluctuations were linked to less efficient early selection mainly during the high tonic-pupil phase in Experiment 1, when frequent-region suppression was still developing.

#### Experiment 2

In Experiment 2, residual pupil size showed little systematic influence on either first fixations on the distractor or first fixations on the target in frequent-region distractor trials. Across tonic-pupil phases, all distractor-fixation credible intervals included zero (largest Pr(β > 0) = .826). The complementary target-fixation model showed the same pattern (largest Pr(β < 0) = .695).

Response type (Go vs. No-Go) differed across experiments and sometimes modulated pre-stimulus residual pupil size (see **Figure 3e and f**), raising the concern that residual-pupil effects might partly reflect response-control demands or carryover rather than state fluctuations. To address this, we ran Bayesian response-composition sensitivity checks using Go-only trials and models with current- and previous-trial response type as covariates. These checks preserved the same qualitative pattern. In Experiment 1, the frequent-region/high-phase residual-pupil slope for first fixations on the distractor remained positive in the Go-only model, β = 0.191, 95% CrI [0.045, 0.349], Pr(β > 0) = .995, and in the response-covariate model, β = 0.181, 95% CrI [0.051, 0.312], Pr(β > 0) = .996. The complementary target-fixation slopes were negative, in the Go-only model, β = -0.127, 95% CrI [-0.232, -0.019], Pr(β < 0) = .989, and in the response-covariate model, β = -0.096, 95% CrI [-0.194, -0.000], Pr(β < 0) = .976. In Experiment 2, no comparable frequent-region fixation-probability effect was observed for first fixations on the distractor or target. Thus, the fixation pattern is unlikely to be explained solely by Go/No-Go composition or immediate response-history carryover.

Together, the first fixation analyses support a pupil-linked learning-phase account anchored primarily in the Experiment 1 effect on first fixations on the distractor. The primary distractor-fixation result shows that residual pupil fluctuations were associated with greater oculomotor capture during the high tonic-pupil phase of Experiment 1, when frequent-region suppression was still developing. The complementary target-fixation result points in the same direction. Later phases of Experiment 1 and the Experiment 2 task context showed less residual-pupil modulation of capture.

### Complementary Fixation-Stage Analysis

Because each trial required participants to inspect two targets, we separated distractor fixations into three stages: before the first target was fixated, between the first and second target fixations, and after both targets had been inspected. The pupil-related fixation-stage results were more selective. Residual-pupil effects were clearest for early distractor fixations in Experiment 1. In the frequent distractor region, larger residual pupil size increased the log-odds of a distractor fixation before the first target in the intermediate tonic-pupil phase (β = 0.202, 95% CrI [0.030, 0.375], Pr(β > 0) = .988), and the rare-minus-frequent residual-pupil slope difference in the same cell was credibly negative (β = -0.732, 95% CrI [-1.146, -0.324], Pr(β < 0) > .999). Other fixation-stage residual-pupil slopes were less stable. Thus, this complementary analysis suggests that residual pre-stimulus pupil size modulated early distractor capture rather than late post-target distractor looking.

### Residual Pupil Effects on Saccade Latency

The Bayesian latency models showed that higher residual pre-stimulus pupil size generally shortened the latency of first saccades that landed on a target (**Figure 5**). In Experiment 1, this target-landing speeding was reliable on frequent-region distractor trials across all tonic-pupil phases (βs = -11.83 to -17.95 ms, all Pr(β < 0) ≥ .987), with additional effects on rare-region distractor trials in the intermediate phase and on distractor-absent trials in the intermediate and low phases. Effects on first saccades that landed on a distractor were narrower: in Experiment 1, higher residual pupil size shortened distractor-landing latencies for frequent-region distractors across phases (βs = -18.47 to -20.51 ms, all Pr(β < 0) ≥ .981), but not reliably for rare-region distractors. In Experiment 2, target-landing speeding was again most evident on frequent-region distractor trials during the high and low tonic-pupil phases (βs ≈ -24 ms, Pr(β < 0) = .998), with additional effects on rare-region distractor trials in the intermediate and low phases. In contrast, Experiment 2 showed no robust residual-pupil effects on distractor-landing latencies. These results indicate that higher trial-level alertness generally facilitates target selection, with the very first saccade being directed straight to a target. Of note, though, while trial-level alertness boosts generally expedite task execution, this may come at the expense of increased performance errors (as revealed by more intricate inter-trial analyses reported in Supplementary “Trial-Level Alertness Fluctuations: Inter-trial Effects”)—indicative of perturbed task control. Thus, whether or not trial-level alertness boosts benefit both the speed and accuracy of processing appears to depend on the operation—fluid versus perturbed—of the task set.

**Figure 5.**
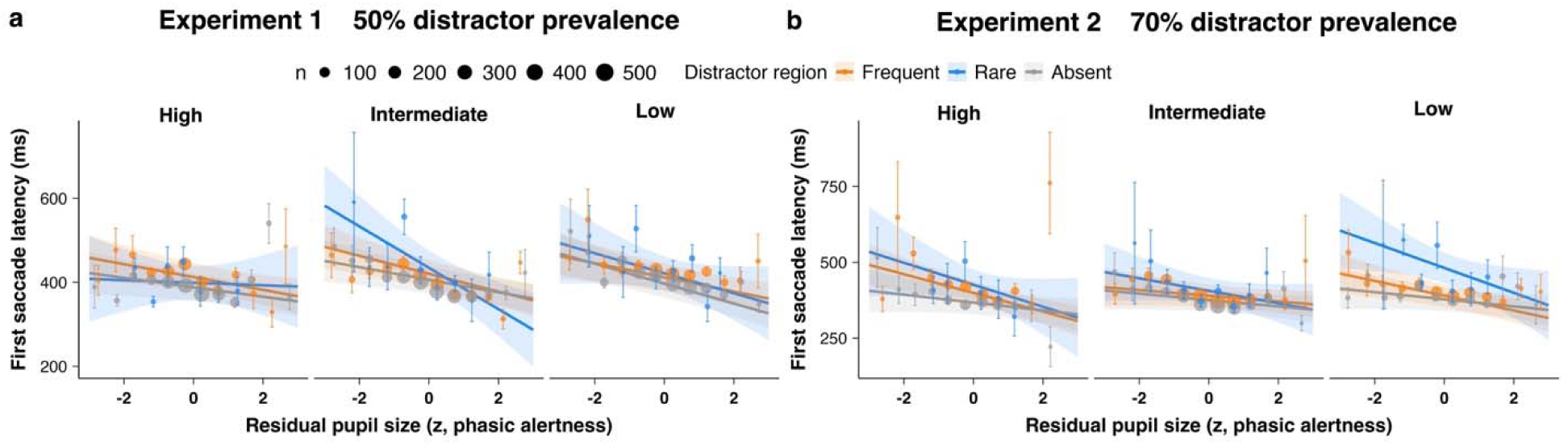
Model-predicted first-saccade latencies as a function of residual pre-stimulus pupil size, tonic-pupil phase, distractor region, and experiment. Solid lines depict model-predicted trends; shaded areas depict 95% credible intervals; points represent binned raw data scaled by trial count (error bars indicate ±1 SE). Lower latency indicates faster target-directed saccades.

## Discussion

We investigated whether relatively enduring, tonic alertness and moment-to-moment, trial-level alertness fluctuations influence how people learn to suppress distractors. Eye-movement behavior revealed robust statistical learning: under moderate distractor prevalence (50%), suppression of frequent-region distractors developed gradually, whereas under high prevalence (70%), it emerged early, within the first block. A formal cross-experiment comparison confirmed this difference, demonstrating that distractor prevalence modulates the temporal trajectory of suppression learning. Next, we examined the vigilance-related mechanisms shaping this learning process, decomposing baseline pupil size into tonic alertness trends and trial-level fluctuations. Pupilometry showed that tonic alertness declined over time and showed a U-shaped relation to RTs, while trial-level fluctuations had distinct, task-dependent effects. With distractors present in the frequent region, trial-to-trial fluctuations in alertness had the greatest impact when suppression was still largely reactive at the initially high level of tonic alertness (in Experiment 1): under these conditions, increased trial-level alertness amplified distractor capture. Once proactive suppression stabilized—reflected in a declining capture rate with reducing levels of tonic alertness—trial-level fluctuations exerted little influence. This transition occurred faster under high distractor prevalence (Experiment 2), reflecting faster adaptation and greater resilience. In contrast, rare-region distractors consistently evoked reactive control, with no systematic modulation by tonic and trial-level alertness.

Decomposing baseline pupil size into tonic trends and trial-level fluctuations revealed a U-shaped relation between tonic alertness and RTs, consistent with the Yerkes-Dodson principle and broadly compatible with the adaptive-gain framework (Aston-Jones & Cohen, 2005; Joshi & Gold, 2020). Importantly, this quadratic relationship emerged specifically for reaction times. For oculomotor capture measures, the tonic alertness–performance relationship appeared more monotonic: intermediate and low tonic states consistently produced fewer distractor fixations and more target fixations than high tonic states, without the performance costs at very low arousal —within the range of arousal states observed here— that would characterize a full inverted-U. This dissociation may reflect the different processing demands underlying each measure. The inverted-U for RT likely captures a Yerkes-Dodson-like optimum for response speed, where both hyper- and hypo-arousal impair the efficiency of response selection and execution. In contrast, oculomotor suppression—which depends on the integrity of proactive spatial suppression—may operate more effectively at stabilized, moderate-to-low arousal levels, where consolidated task sets support consistent suppression without the interference from heightened trial-level responsivity that accompanies high tonic arousal. This was echoed in the first-fixation data: intermediate or low tonic states led to fewer saccades to distractors and more target fixations, reflecting effective suppression and goal-directed processing (see also Sara, 2009; Unsworth & Robison, 2016). Given the gradual decline in baseline pupil size across trial blocks, indicative of a reduction in tonic alertness, this optimal performance pattern may reflect a stabilized state of distractor suppression. Of note, over the course of the lengthy experiments, tonic alertness declined asymptotically towards a level still supporting efficient goal-directed processing, rather than dropping to the very low, disengaged state characteristic of hypo-activity (Aston-Jones & Cohen, 2005).

Trial-level alertness fluctuations in pupil size, in turn, aligned with predictive-coding theories of attention, which view attentional adjustments as responses to prediction errors—signals of deviation from expectation—that update attentional states in real time (Clark, 2013; Friston, 2005). In Experiment 1, the rare-region distractors (10% probability) produced the largest alertness increases, indicating that their unexpected occurrence triggered greater surprise. In turn, these larger trial-level responses predicted faster RTs, suggesting that rare-region distractors on the preceding trial n–1 produced an “alerting signal” that increased general system ‘readiness’ on the current trial n (Posner & Petersen, 1990; Petersen et al., 2017). However, this was associated with an increase in error rates (see Supplementary “Trial-Level Alertness Fluctuations: Inter-trial Effects”)—suggestive of a momentary perturbation of task control. Thus, whether or not trial-level alertness boosts are purely beneficial to performance appears to depend on the task set remaining intact versus being perturbed by the previous distractor event. In Experiment 2, trial-level alertness was linked to response type rather than distractor region: under higher distractor prevalence (70%), shorter display durations, and greater Go/No-Go uncertainty, the Go/No-Go decision on trial n–1 carried greater behavioral weight and became the primary driver of alertness fluctuations carried over to trial n. Together, these findings indicate that, rather than being a random or purely spontaneous state fluctuation, trial-level alertness is responsive to the context and moment-to-moment demands of the task.

Besides distinct but complementary influences of tonic and trial-level alertness on behavior, most importantly, we found that these two forms of alertness shape distractor suppression by modulating the reliance on proactive versus reactive control. When distractor prevalence was moderate (50%, Experiment 1), suppression was initially dominated by reactive mechanisms, requiring substantial cognitive effort to stabilize the task set, reflected in elevated tonic alertness. During this phase, performance was highly sensitive to trial-level alertness: higher trial-level alertness increased distractor capture, a hallmark of reactive control (Braver, 2012). As learning progressed and proactive suppression strengthened, trial-level alertness exerted a diminishing influence on distractor capture. When distractor prevalence was high (70%, Experiment 2), suppression shifted rapidly to a proactive mode, leaving trial-level alertness minimal impact on distractor handling. Once consolidated, proactive suppression operated automatically, relatively impervious to trial-level fluctuations. Seemingly at variance with the detrimental impact of trial-level alertness increases on oculomotor capture (by frequent-region distractors at high tonic alertness), such increases generally (independently of distractor condition and tonic alertness) expedited the generation of first saccades directly to a target. This suggests that, on trials in which the task set is implemented optimally, trial-level alertness increases enhance goal-oriented processing (operating fluently without interference).

These findings support a framework (**Figure 6**) in which task statistics and tonic/trial-level alertness interact dynamically. Global distractor prevalence and local distractor distributions determine whether suppression must operate reactively or can be applied proactively through learned routines. Early in the task, when goal-oriented processing is being established and distractor occurrence is unpredictable, suppression is largely reactive, and higher trial-level alertness may paradoxically increase distractor capture. With practice, tonic alertness decreases to a lower-energy level while proactive suppression develops, becoming efficient and largely insensitive to trial-level fluctuations. Nevertheless, trial-level alertness continues to exert influence through internally generated ‘alerting signals’ (cf. Posner et al., 1973), enhancing focus and facilitating performance on subsequent trials. In sum, tonic alertness dynamically modulates trial-level effects, shaping the balance between reactive and proactive suppression and influencing both goal-related target selection and inadvertent distractor capture. More distally, this framework may also inform neuroergonomic and psychophysiological work by suggesting that pupillometric measures can be used to separate slower state changes from trial-level fluctuations when modeling adaptive attentional control over extended task performance.

**Figure 6.**
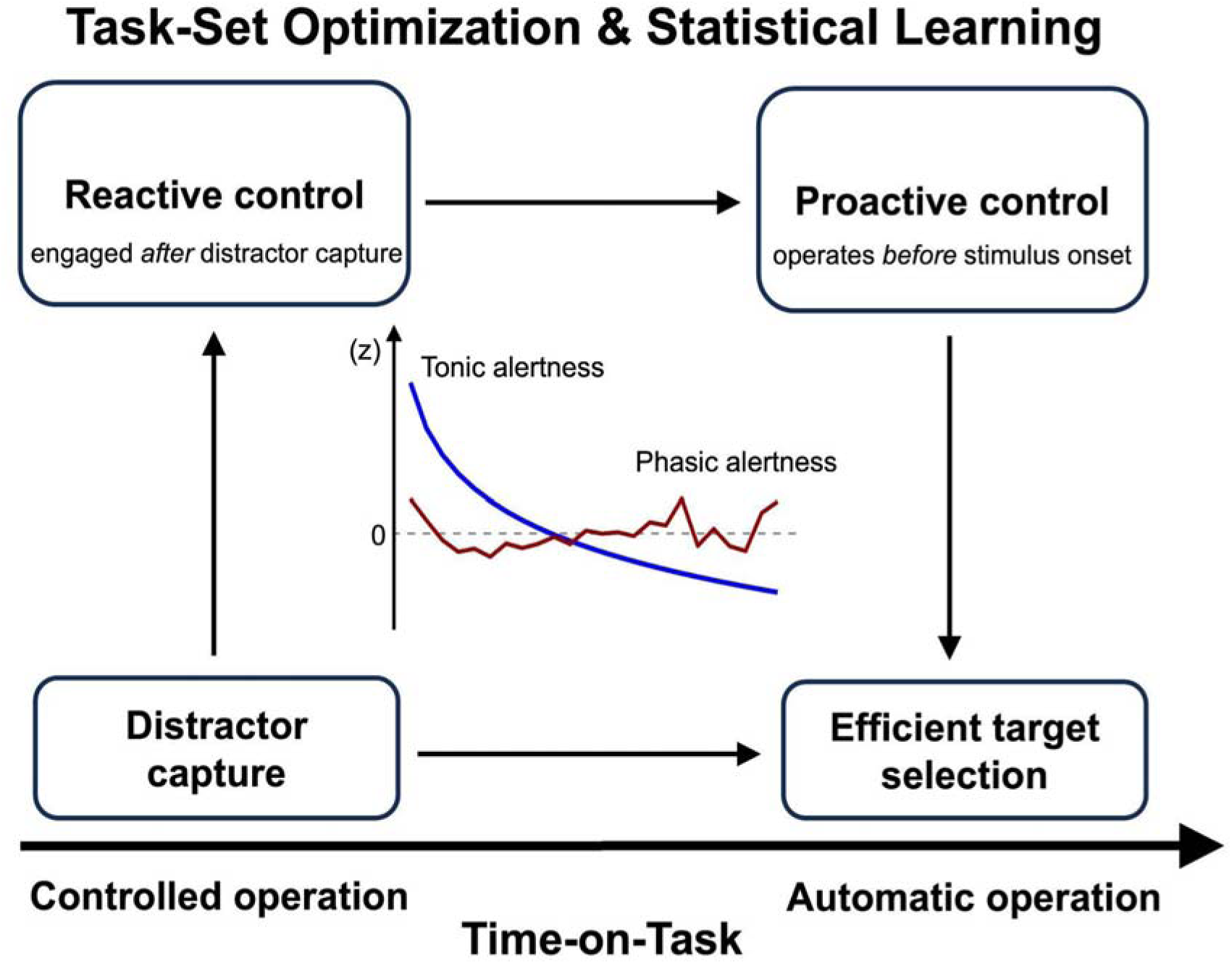
Conceptual framework of dynamic alertness modulation. Attentional control evolves from the interplay between the task set, statistical learning, and tonic and trial-level alertness fluctuations. In the initial set-up and running-in stage, involving high tonic alertness, the task set encompasses the search- and response-critical target features and the S–R mapping. At this stage, distractor control is largely reactive, responding to capture incidents. With increasing experience, associated with a reduction in tonic alertness, the distractor statistics—global prevalence and local distribution—are being acquired and incorporated into the task set, supporting proactive control. Thus, statistical learning drives the transition from reactive to proactive control, while tonic alertness modulates how strongly trial-level alertness signals influence the moment-to-moment incidence of distractor capture and efficiency of target selection.

## Acknowledgments

This work was supported by the German Research Foundation (DFG) grants CH 3093/1-1 and SH 166/10-1, awarded to SC and ZS, respectively. We thank Fredrik Allenmark and colleagues for sharing the raw eye-tracking data, which allowed us to extract and analyze the pupil-size measures reported here.

## Supplemental Materials

### Tonic and Phasic Pupil Dynamics in Error Responses

Errors (binary: 0 = correct, 1 = error) were analyzed using a Bayesian logistic mixed-effects model. The model included tonic alertness (measured by the standardized baseline pupil size, with linear and quadratic terms), phasic vigilance (residual pupil size), distractor region (distractor-absent, frequent, rare), response type (go, no-go), and their interactions, as well as log(block), with random intercepts and slopes for participants.

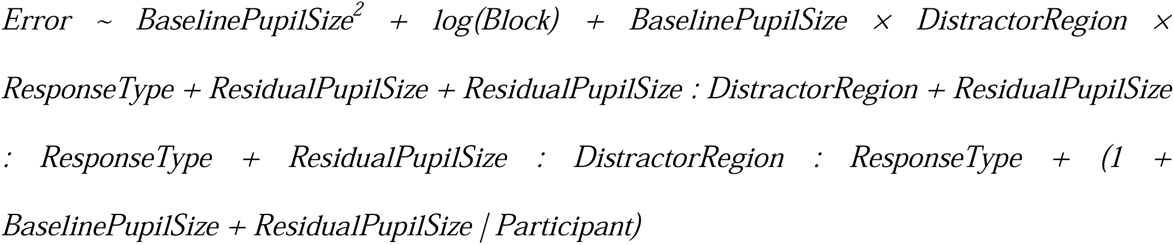

In Experiment 1 (see **Figure S1**), a weak positive quadratic trend for baseline pupil size was observed (0.06, [0.00, 0.12]). Several interactions involving pupil-linked measures were notable. Baseline pupil size interacted with response type: larger baseline pupil size predicted higher error rates on Go versus No-go trials (0.47, [0.08, 0.88]). Baseline pupil size also positively interacted with distractor region, with an increased slope for rare-region versus frequent-region (Rare-Frequent: 0.88, [0.08, 0.88]) and distractor-absent conditions (Rare-Absent: 0.77, [0.22, 1.3]). Higher residual pupil size significantly predicted fewer errors for Go trials (-0.58, [-1.08, -0.11]), while this effect was non-significant for No-Go trials. Higher residual pupil size also predicted reduced errors for rare-region distractor trials (–1.00, [–1.80, -0.14]), but not for frequent-region distractor and distractor-absent trials. In Experiment 2, the overall error was also rather low (–2.75, [–3.50, –2.00]), and it decreased reliably across blocks (–0.29 [–0.46, –0.11]). Baseline pupil size showed a slight, but uncertain, positive association with errors. Phasic increases in pupil size significantly reduced errors on frequent-region distractor trials (–0.63 [–1.07, –0.17]) but not on trials with rare-region or absent distractors. Also, they significantly reduced errors on Go trials (–0.72 [–1.21, –0.24]), which required an overt 2AFC response decision, but not on No-go trials, on which responding was to be withheld. In Experiment 2, the reduced proportion of trials requiring an overt response (60% vs. 80% No-go trials in Experiment 1), coupled with the increased distractor prevalence (70% vs. 50%), which made rare-region distractors more frequent overall, may have rendered error rates more sensitive to residual pupil size than reaction times. These findings suggest that phasic alertness contributes to adaptive performance, though with its influence (on response speed or accuracy) being condition-dependent.

In both Experiments 1 and 2, substantial variability was observed at the participant level in intercepts (both experiments, SD > 0.76) and in slopes for baseline (both, SD > 0.44) and residual pupil size (both, SD > 0.41). Notably, the slopes for baseline and residual pupil size were strongly negatively correlated in the two experiments (both r = –0.87), indicating that participants who showed stronger tonic-alertness effects tended to exhibit weaker phasic modulation, and vice versa. This pattern suggests that tonic and phasic components of alertness represent distinct yet complementary mechanisms supporting attentional control.

**Figure S1.**
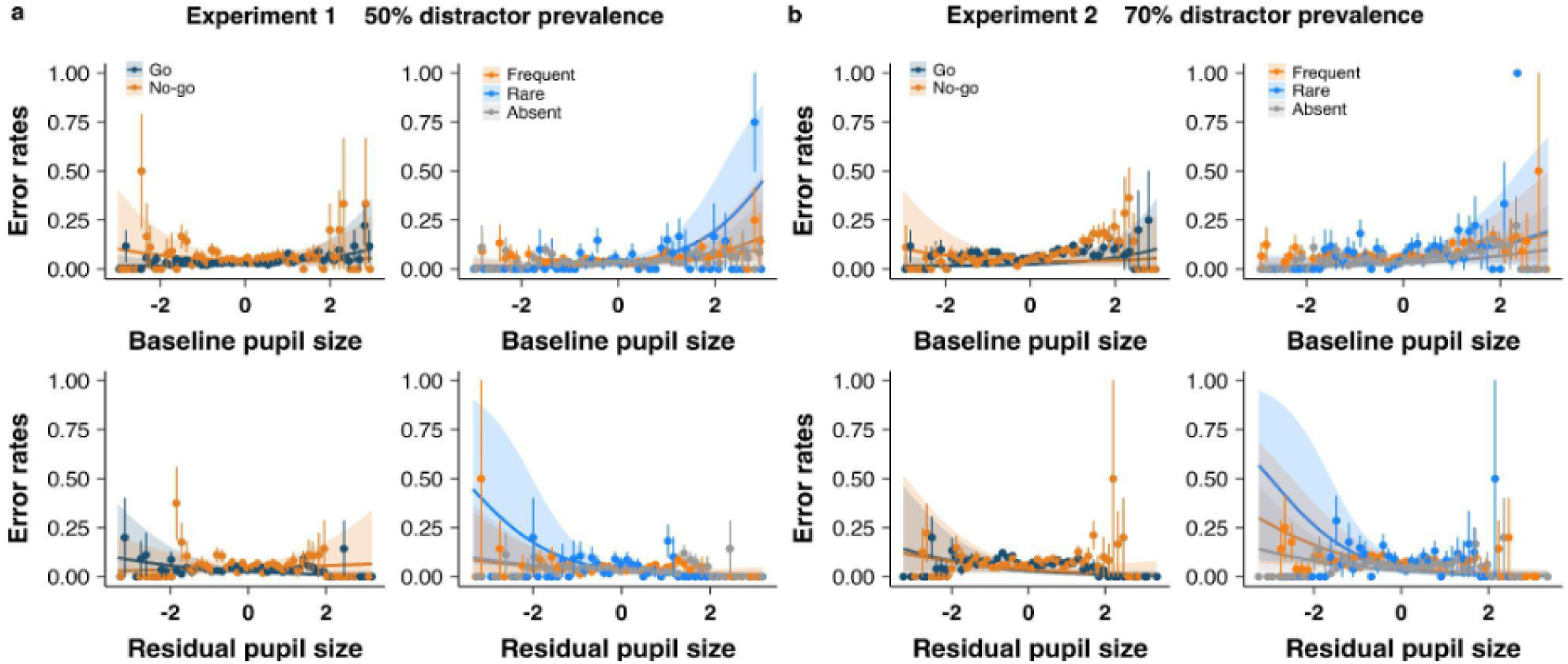
**a** and **b** depict the effects of tonic and phasic pupil-size measures on error rates as a function of response type (Go, No-go) and distractor region (Frequent, Rare, Absent) in Experiments 1 and 2, respectively. Points represent binned averages (30 bins per distractor region), and lines depict model-predicted quadratic/linear fits for each distractor-region or response-type condition. Shaded ribbons = 95% CI. Error bar = SE.

### Trial-Level Alertness Fluctuations: Inter-trial Effects

In Experiment 1, when a rare-region distractor captured the eye on the previous trial *n–1*, performance was more prone to errors on the current trial *n* (*F*(1,14) = 8.26, *p* = .012, *η_p_*^2^= .37), coupled with shorter reaction times on correct trials (*F*(1,14) = 9.53, *p* = .008, *η_p_*^2^= .41), compared to when the eye was directed straight to a target, avoiding the rare-region distractor, on the trial *n–1* (error rates: 4% vs.1%; RTs: 1061 ms vs. 1154 ms; see **Figure S2**). No such effects on the current trial were found when there was a distractor in the frequent region on the preceding trial (ps > .15, ηₚ²s < .14). In Experiment 2, responses on a given trial *n* were only non-reliably faster when a previous rare-region distractor had captured the eye compared to when a target was the first item fixated (1135 ms vs. 1202 ms; *p* = .37, *ηₚ²* = .06), with little difference in the (compared to Experiment 1 overall increased) error rates (6% vs. 8%; *p* = .20, *ηₚ²* = .10).

Of note, for Experiment 1, a 2 (Distractor Region: rare vs. frequent) x 2 (First Fixation: on distractor vs. on target) ANOVA of *phasic alertness induced on trial n–1* revealed only a main effect of Distractor Region (*F*(1, 14) = 6.62, *p* = .022, *ηₚ²* = .32), but no effects involving First Fixation (main effect: *F*(1, 14) = 0.02, *p* = .883, *ηₚ²* < .01; interaction: *F*(1, 14) = 0.62, *p* = .446, *ηₚ²* = .04). Thus, the phasic alertness increases induced by rare-region distractors were comparable whether or not the distractor captured the eye. Yet, a subsequent speeding of responses coupled with an increase in error rates was observed only when the (rare-region) distractor caused oculomotor capture. This dissociation suggests that phasic alertness springs from *pre-attentive* registration of an unexpected distractor event—eliciting high surprise—as such, whereas behavioral adjustments on the subsequent trial critically depend on the extra control demands arising when the distractor actually captured the gaze.

In summary, these findings suggest that, while phasic alertness induced by a rare-region distractor that captured the eye on trial *n–1* speeds up processing on the current trial *n*, this comes at the expense of an increase in error rates. That is, attentional capture by rare-region distractors causes a momentary lapse in speed-accuracy trade-off monitoring, indicative of perturbed task control. No such perturbation was evident when frequent-region distractors captured the eye: such distractors appear to be handled in a rather efficient, ‘routinized’ fashion (see, e.g., Sauter et al., 2021, who observed greatly shortened fixational dwell times on frequent-region, compared to rare-region, distractors). Importantly, this effect pattern emerged when distractor prevalence was moderate (50%), but not when distractors occurred on the majority of trials (70%)—suggesting that with higher distractor prevalence, perturbations of task control by high-surprise events become less evident.

**Figure S2.**
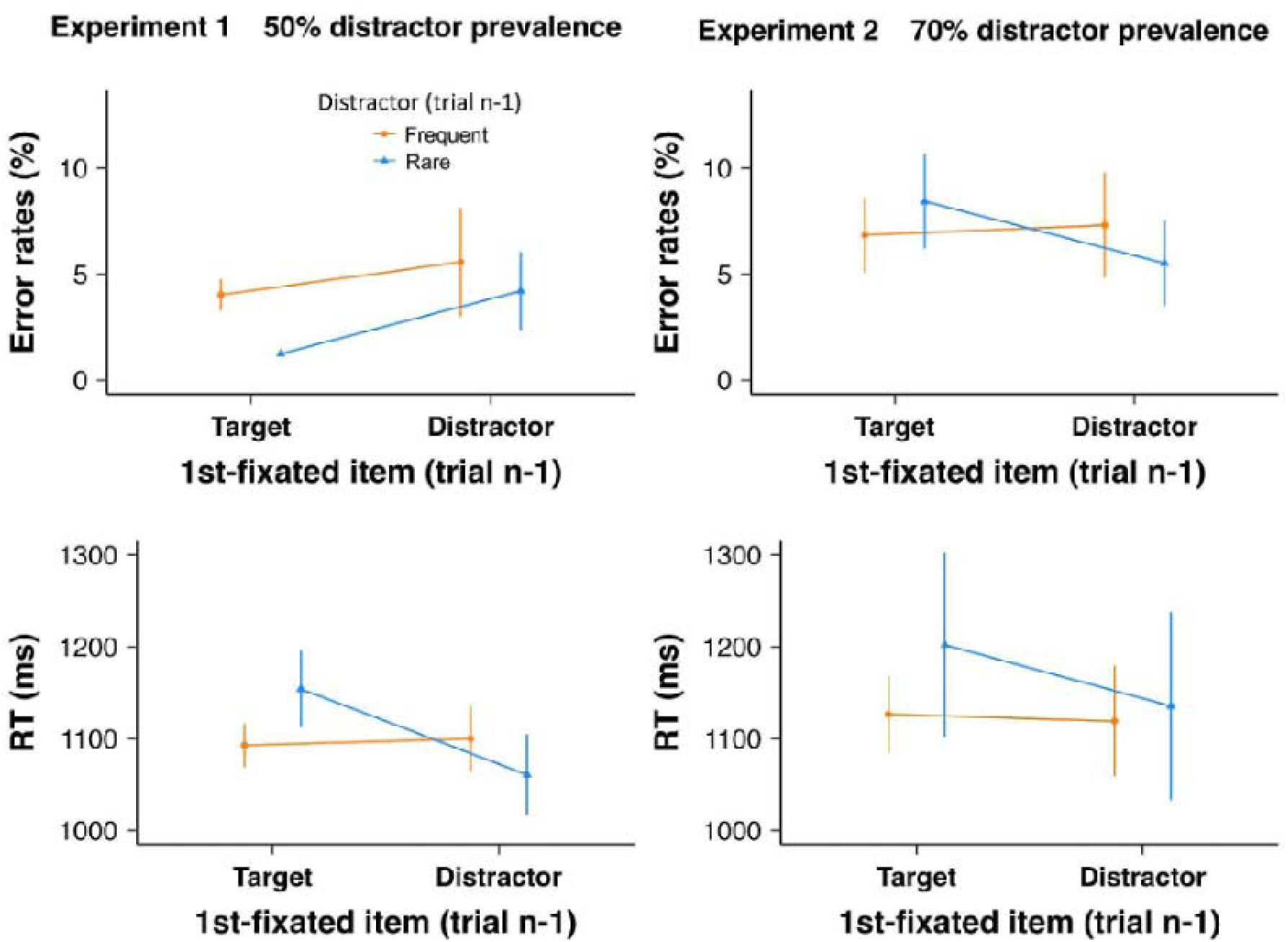
Error rates (%) and RTs (ms) on a given trial n as a function of the distractor region (Frequent, Rare) and the item fixated first (Distractor, Target) on the preceding trial n–1, separately for Experiment 1 (**left**) and Experiment 2 (**right**). Error bars indicate ±1 SE.

### Stimulus-Triggered Pupil Response Analysis

In both experiments, the stimulus-triggered pupil response showed a characteristic pattern across all distractor region conditions: a brief dilation (0–400 ms post-stimulus), followed by a pronounced constriction and subsequent recovery (see **Figure S3**). The constriction peaked at approximately 690 ms in Experiment 1 (N = 15; 6,682 frequent, 840 rare, 7,564 absent trials) with an amplitude of ∼14%, and at ∼610 ms in Experiment 2 (N = 16; 9,895 frequent, 1,059 rare, 5,539 absent trials) with a smaller amplitude of ∼7%. Critically, the factor distractor region had no significant effect on the pupil time course in either experiment. Cluster-based permutation tests (10,000 permutations) revealed no significant clusters—in fact, no time points exceeded the cluster-forming threshold of p < .05—either collapsed across response type or separately for Go and No-go trials (see **Figure S4**). Confirmatory window-based ANOVAs on participant-level means likewise showed no effects: all Fs < 0.10, all ps > .90 across peak (500–700 ms), sustained (700–1,000 ms), and late (1,000–1,200 ms) windows in both experiments. Descriptively, No-go trials in Experiment 2 showed a trend toward reduced constriction for rare-region distractors (M = −1.6%) relative to frequent-region (M = −2.6%) and absent (M = −1.7%) conditions in the sustained window (see **Figure S4, panel D**), but this did not approach the cluster-forming threshold. With N = 15–16, the cluster-based permutation test had 80% power to detect effects of d_z_ ≥ 0.75–0.78 (large). But the observed F-values were essentially zero, so the true effect—if any—is negligible, not merely undetected. The null result proves that observed pupil effects in the main texts reflect baseline arousal states—not sensory responses to luminance or display configurations. The temporal dissociation between the lack of within-trial effects and the presence of subsequent-trial baseline shifts suggests that distractor-driven alertness unfolds slowly over the inter-trial interval.

**Figure S3.**
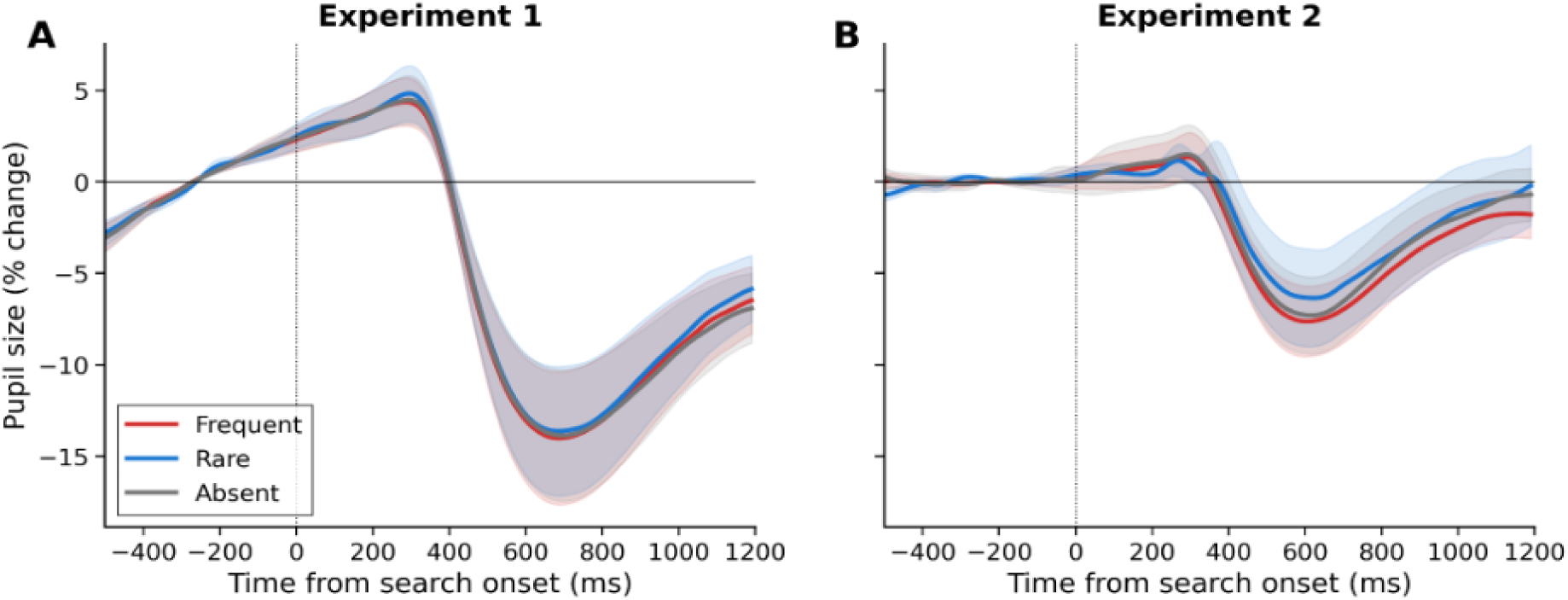
Stimulus-triggered pupil response by distractor region (Go + No-go collapsed). Shaded areas represent ±1 SE across participants. Dotted line indicates search display onset.

**Figure S4.**
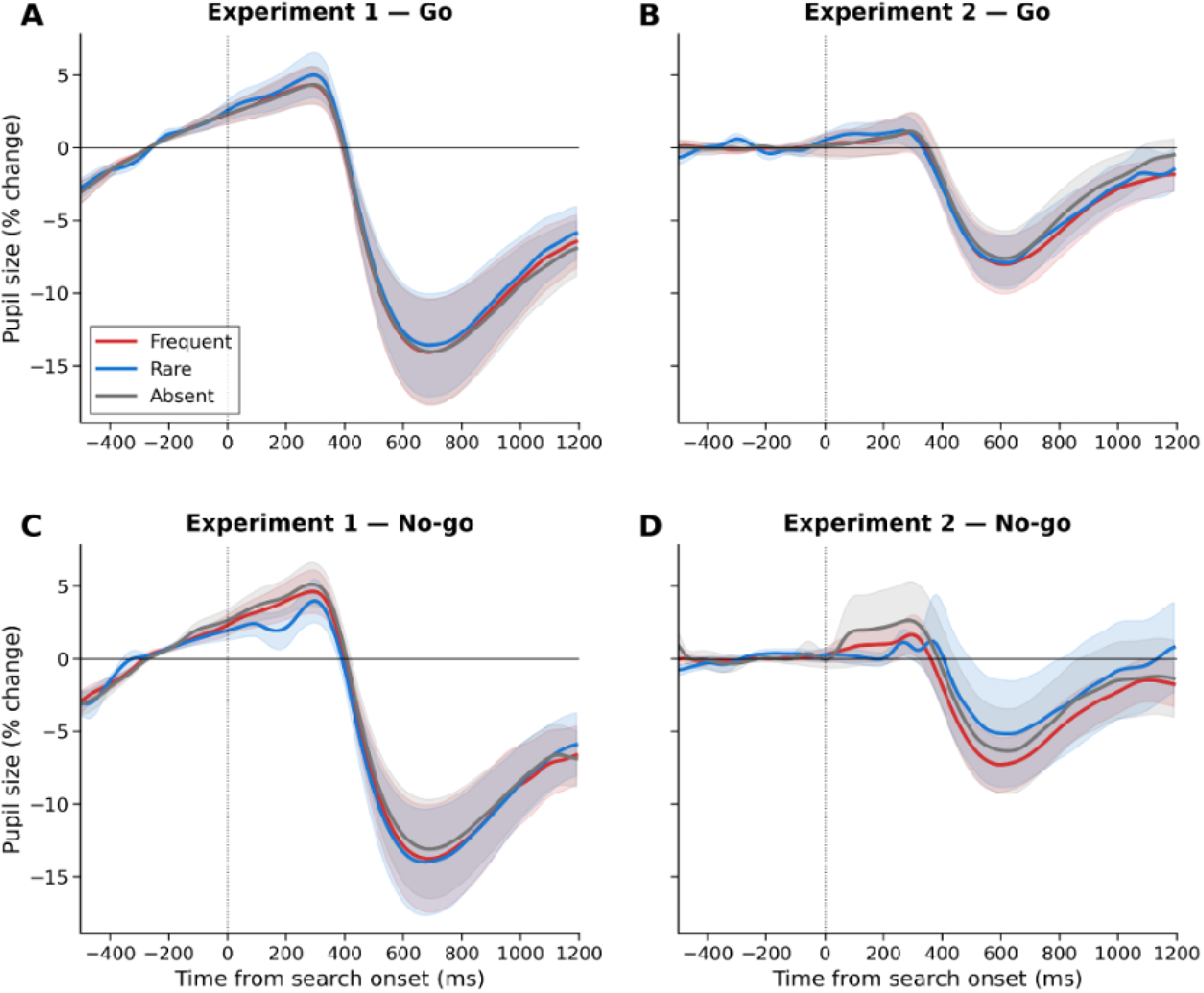
Stimulus-triggered pupil response by distractor region, separately for Go (top) and No-go (bottom) trials. Shaded areas represent ±1 SE across participants.

