## Supplementary for "Dynamic Modulation of Distractor Suppression by Tonic and Trial-Level Alertness Fluctuations: A Pupillometric Study"

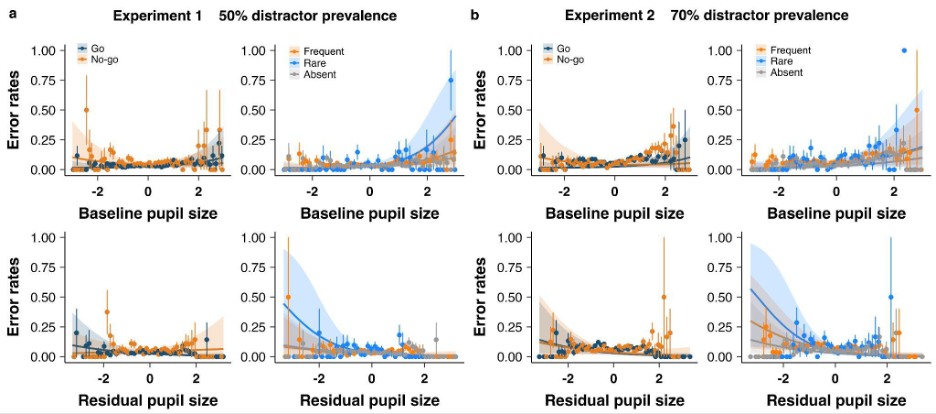

**Figure S1.** ***a*** *and* ***b*** *depict the effects of tonic and phasic pupil-size measures on error rates as a function of response type (Go, No-go) and distractor region (Frequent, Rare, Absent) in Experiments 1 and 2, respectively. Points represent binned averages (30 bins per distractor region), and lines depict model-predicted quadratic/linear fits for each distractor-region or response-type condition. Shaded ribbons = 95% CI. Error bar = SEM.*

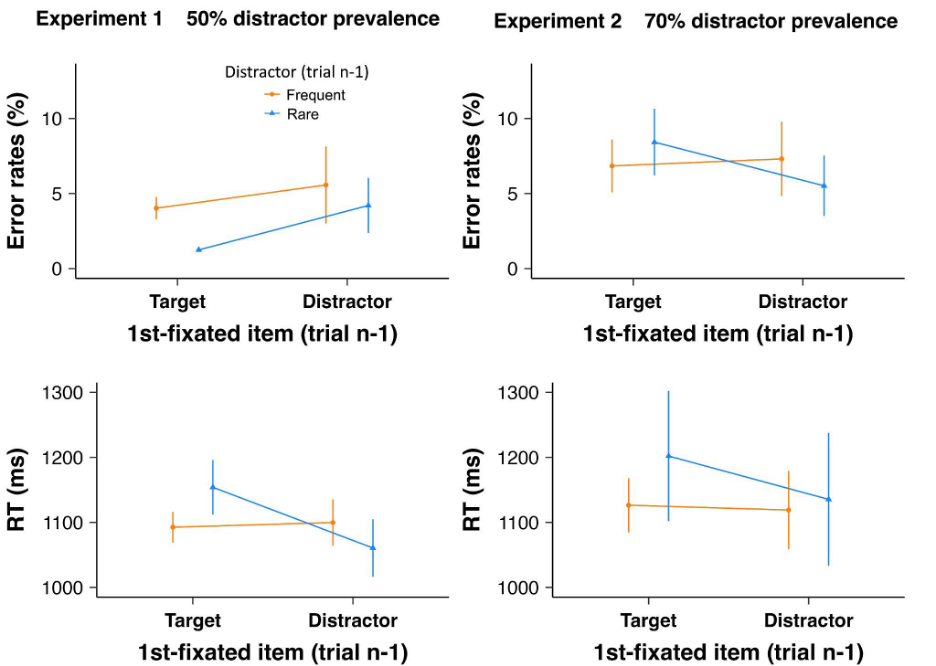

***Figure S2.*** *Error rates (%) and RTs (ms) on a given trial n as a function of the distractor region (Frequent, Rare) and the item fixated first (Distractor, Target) on the preceding trial n–1, separately for Experiment 1 (****left****) and Experiment 2 (****right****). Error bars indicate ±1 SE.*

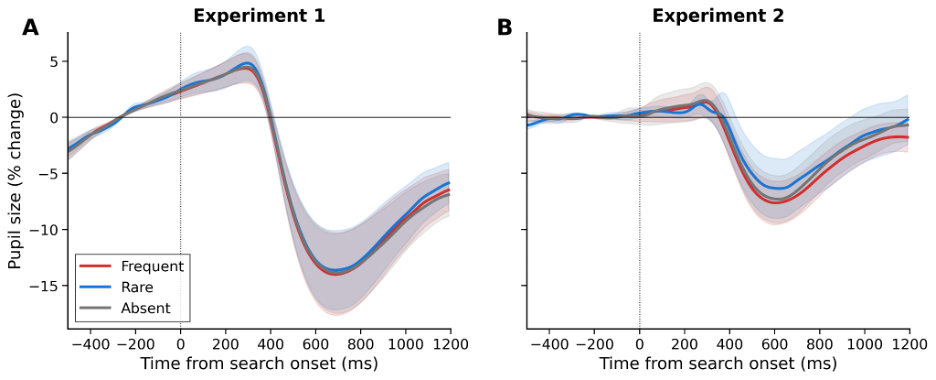

***Figure S3.*** *Stimulus-triggered pupil response by distractor region (Go + No-go collapsed). Shaded areas represent ±1 SE across participants. Dotted line indicates search display onset.*

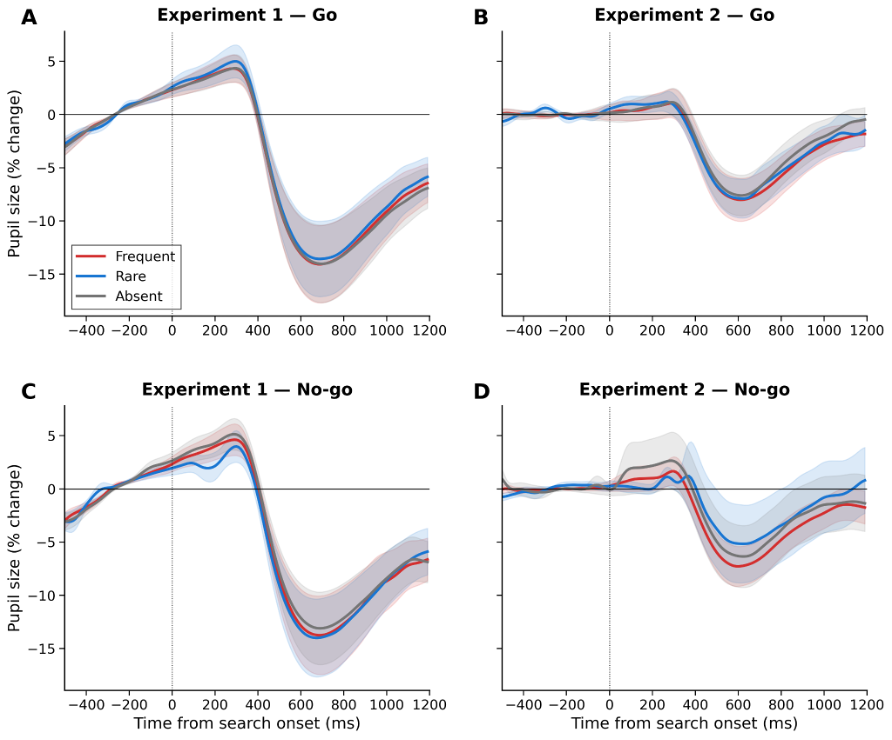

***Figure S4****. Stimulus-triggered pupil response by distractor region, separately for Go (top) and No-go (bottom) trials. Shaded areas represent ±1 SE across participants.*
